# A phenomic modeling approach for using chlorophyll-a fluorescence-based measurements on coral photosymbionts: a step towards bio-optical bleaching prediction

**DOI:** 10.1101/2022.11.07.515332

**Authors:** Kenneth D. Hoadley, Grant Lockridge, Audrey McQuagge, K. Blue Pahl, Sean Lowry, Sophie Wong, Zachary Craig, Chelsea Petrik, Courtney Klepac, Erinn M. Muller

## Abstract

We test a newly developed instrument prototype which utilizes time-resolved chlorophyll-*a* fluorescence techniques and fluctuating light to characterize Symbiodiniaceae functional traits across seven different coral species under cultivation as part of ongoing restoration efforts in the Florida Keys. While traditional chlorophyll-*a* fluorescence techniques only provide a handful of algal biometrics, the system and protocol we have developed generates > 1000 dynamic measurements in a short (∼11 min) time frame. Resulting ‘high-content’ algal biometric data revealed distinct phenotypes, which broadly corresponded to clade-level Symbiodiniaceae designations determined using quantitative PCR. Next, algal biometric data from *Acropora cervicornis* (10 genotypes) and *A. palmata* (5 genotypes) coral fragments was correlated with bleaching response metrics collected after a two month-long exposure to high temperature. A network analysis identified 1973 correlations (Spearman R > 0.5) between algal biometrics and various bleaching response metrics. These identified biomarkers of thermal stress were then utilized to train a predictive model, and when tested against the same *A. cervicornis* and *A. palmata* coral fragments, yielded high correlation (R = 0.92) with measured thermal response (reductions in absorbance by chlorophyll-a). When applied to all seven coral species, the model ranked fragments dominated by *Cladocopium* or *Breviolum* symbionts as more bleaching susceptible than corals harboring thermally tolerant symbionts (*Durusdinium*). While direct testing of bleaching predictions on novel genotypes is still needed, our device and modeling pipeline may help broaden the scalability of existing approaches for determining thermal tolerance in reef corals. Our instrument prototype and analytical pipeline aligns with recent coral restoration assessments that call for the development of novel tools for improving scalability of coral restoration programs.

## Introduction

Rising ocean temperature due to climate change is the greatest global threat to coral reefs (Hughes et al., 2017; Hughes et al., 2018). Traditional reef management strategies can be effective at mitigating local stressors (e.g., urban encroachment, pollution, and overfishing) (Gattuso et al., 2018; Donovan et al., 2021) but will not prevent rapid coral loss caused by increasingly frequent mass bleaching events. Active coral conservation programs that focus on direct restoration are needed and offer a means to mitigate the effects of global warming while more permanent solutions for combating climate change are developed (Boström-Einarsson et al., 2020; Caruso et al., 2021; Voolstra et al., 2021a). Indeed, reef management programs are increasingly reliant on active restoration activities including the cultivation and transplant of coral fragments to improve community resilience and/or restore impacted reefs (assisted evolution; (Van Oppen et al., 2015; Baums et al., 2019; Rinkevich, 2019; Grummer et al., 2022). Ideally, transplanted corals should have positive heritable traits that increase thermal resilience among the coral population, thereby minimizing their susceptibility to future thermal anomalies. Reliable and scalable identification of thermally tolerant coral genotypes is thus a critical need for improved efficacy in certain coral restoration initiatives (Voolstra et al., 2021a; Grummer et al., 2022). Recent studies that utilize molecular techniques to investigate linkage between thermal tolerance and coral host genetics (Fuller et al., 2020; Drury & Lirman, 2021; Drury et al., 2022) or microbial community dynamics (Peixoto et al., 2017; Ziegler et al., 2017; Rosado et al., 2019) are promising, but not yet fully developed. Others that rely on optical properties of the symbiotic algae (Suggett et al., 2022) also hold significant promise but need to be further contextualized for broader accessibility and scale.

The dinoflagellate family Symbiodiniaceae is a genetically diverse group of photosynthetic alga, many of which are important primary producers and often found in symbioses with reef-building corals (LaJeunesse et al., 2018). Tolerance to environmental perturbations such as high temperature events differ across Symbiodiniaceae species (Suggett et al., 2017; LaJeunesse et al., 2018), significantly impacting a coral’s overall susceptibility to thermal bleaching events (Hughes et al., 2017). Distinguishing Symbiodiniaceae species has been almost exclusively done through genetic testing and prior knowledge of the species’ capacity to cope with high temperature events. Certain photosynthetic traits differ across symbiont types or during exposure to thermal stress (Suggett et al., 2015; Hoadley et al., 2021), providing a suite of potentially useful biomarkers of bleaching susceptibility. Importantly, many of these traits can be rapidly measured using non-destructive bio-optical means.

Within reef corals, photosynthetic breakdown due to thermal stress has been widely described using chlorophyll a fluorescence (here on referred to as CF) techniques which are relatively instantaneous and can provide a wealth of information regarding the capture and use of photons within living photosynthetic organisms (Kolber et al., 1998; Gorbunov et al., 2001; Papageorgiou, 2007). These tools have contributed greatly to our current understanding of thermal tolerance across individual Symbiodiniaceae species (Warner et al., 1999). However, most studies have relied on instrumentation that use relatively long pulses of light (known as multi-turnover instruments such as the Walz PAM) which limits their capability to interrogate highly dynamic CF signatures. More sophisticated instrumentation that utilize short-pulses of light (known as ‘single-turnover’ instruments) to interrogate photochemical parameters, can provide a more dynamic and informative dataset (Kolber et al., 1998; Ragni et al., 2010; Suggett et al., 2015). With certain notable exceptions, CF methods that utilize single-turnover instrumentation can resolve some photo-physiological traits across phylogenetic relationships, demonstrating the techniques’ promise as a tool for the assessment of functional differences across symbiont types (Hennige et al., 2007; Hennige et al., 2009; Suggett et al., 2015; Hoadley et al., 2021). In addition, advancements in our understanding of dynamic and stochastic light conditions suggest that rapid changes in photo-acclimation measured during short time frames also provides valuable information for assessing photo-physiology (Allahverdiyeva & Suorsa…, 2015; Andersson et al., 2019). Methods that incorporate single-turnover measurements to capture photochemical responses to dynamic changes in light could therefore greatly improve our ability to resolve functional differences across Symbiodiniaceae species and provide valuable insight to basic and applied fields of coral reef science.

Here we introduce a newly developed, low-cost, single turnover fluorometer specifically designed for interrogation of symbiotic algae *in hospite*. This instrument was designed on previously established and well-developed principles (Kolber et al., 1998) but also utilizes multiple excitation wavelengths (420, 442, 458, 505, and 520 nm) to preferentially excite different photopigments and light-harvesting compounds which may differ across symbiont types or environmental conditions (Hennige et al., 2009). Our time-resolved protocols compared a suite of spectrally dependent photochemical parameters (Table 1) as they respond to fluctuating light and generate thousands of dynamic photo-physiological measurements during a short 11 min sampling protocol. This technique was then utilized to generate distinct phenotypes across seven different coral species and our results demonstrate how bio-optical approaches can be used to identify important photo-physiological differences across various coral species and light environments. Next, our phenotypic approach was utilized to train a predictive bleaching model by linking photo-physiological metrics with bleaching responses from two coral species (*Acropora palmata* and *Acropora cervicornis*). High accuracy of the predictive model was confirmed against the same coral genets used in a bleaching experiment, demonstrating the cumulative power of using thousands of photo-physiological based biomarkers to assess bleaching susceptibility. A truncated version of the model was then applied to genets from the previously measured seven coral species. Results broadly correspond with underlying symbiont types, indicating good accuracy even across coral species and symbiont types not included in the original training dataset. Tools that improve scalability and accessibility for identifying functional traits such as bleaching susceptibility are valuable to the coral restoration field (Voolstra et al., 2020) and our novel instrumentation and analytical pipeline may benefit certain restoration initiatives.

**Table 1.**
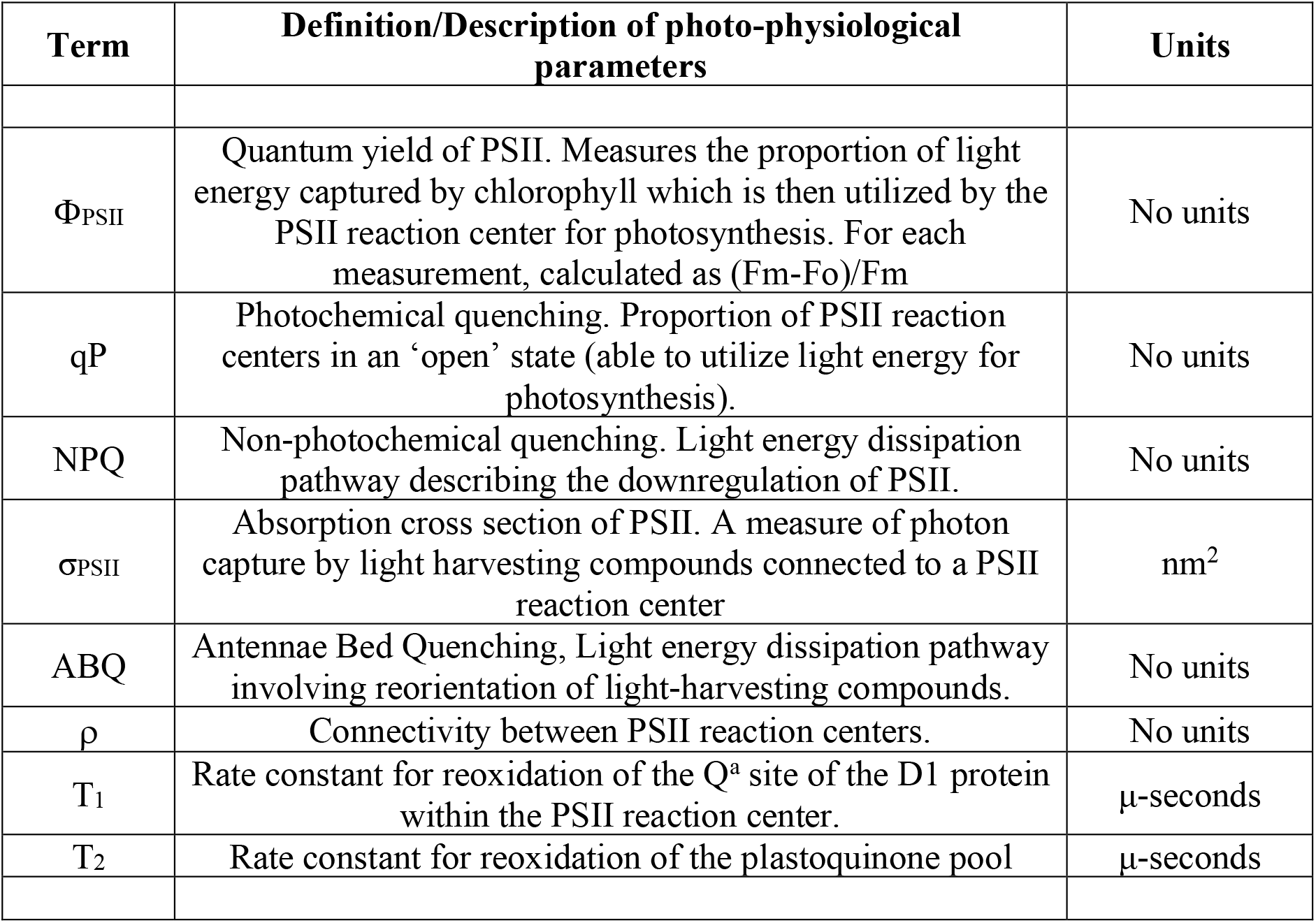
Table of photo-physiological parameters, units, and definitions/descriptions. Within this study, each defined parameter is represented by spectrally dependent values at each measurement time point (see Fig. 1).

## Materials and Methods

### Construction of Multispectral Fluorometer

A detailed list of parts, along with PCB board, housing and schematics can be found in the supplemental materials (fig. S1) and through an online repository (GitHub: khoadley/MultSpectral-ST-PCB). Briefly, the multispectral fluorometer is controlled via a small microcontroller (Teensy 4.1) which governs both the emitter and detector assemblies. Each emitter is comprised of a different color (UV-420nm, Royal Blue-442nm, Blue-458nm, Cyan-505nm, and Green-525nm) 10mm square LED (Luxeon Star) mounted to a 25mm diameter end cap plug (Thorlabs). All LED emitters are coupled using SMA connectors to a custom fiber optic assembly that randomly combines emission and fluorescence detection fibers within a 1-cm diameter probe tip (Berkshire photonics). Maximum LED output is achieved within ∼500 ns using a custom LED driver circuit each of which is individually controlled by the microcontroller. A single white LED (Luxeon Star) is regulated using a 350mA constant current BuckPuck LED driver (LuxDrive) and serves as a variable actinic light source controlled via a ‘PWM to analog’ signal from the microcontroller. A lowpass (<650nm) glass filter was placed between the actinic white LED and fiber optic cable, ensuring minimal stray light can reach the detector. Fluorescence detection is measured using a variable gain avalanche photodiode (Thorlabs). Using a combination of high pass (>650nm) and lowpass (<700nm) glass filters sandwiched between two aspheric lenses (Newport corp), fluorescent light from samples travel from the fiber optic cable and are concentrated onto the detector. Analog signals from the detector are then interpreted using a 12-bit analog to digital converter embedded within the microcontroller. All LED drivers and the detector circuit are housed within a single (13” x 4”) printed circuit board (PCB) (GitHub: khoadley/MultiSpectral-ST-PCB, fig. S1). Fluorescence measurements are stored on an onboard micro-SD card. Firmware for the teensy microcontroller is written using the Arduino IDE and a python-based software program communicates with the microcontroller and manages data capture. All firmware and software are open-source and available via GitHub (khoadley/MultiSpectral-ST-PCB). PCB and electronic components cost between 200-300 USD. Depending on construction materials utilized, complete construction of the fluorometer prototype costs between 4.5-5.5K USD.

Fluorescence measurements consist of an induction curve produced through excitation with 1.3-*µ*s single turnover flashlets spaced apart by 3.4 *µ*s dark intervals (32 flashlets were utilized under 420, 448, and 470-nm excitation while 40 flashlets were utilized during 505 and 520-nm excitation and ensure that all measurements achieved full saturation). After fluorescence induction, a 300 ms relaxation phase consisting of 1.3-*µs* light flashes spaced apart with exponentially increasing dark periods (starting with 59-*µs*). LED power output estimates were determined using the following equation: I/D where I is irradiance measured at a reduced duty cycle (D = 1.3:500) using a 4pi light meter (Walz; µmol m^-2^ sec^-1^). The 1.3 value within the D ratio reflects the length (in *µ*s) of each flashlet and allows us to calculate irradiance while only running the LED at a 1.3:500 duty cycle. Power (irradiance in PAR) estimates were then used to determine the spectrally dependent functional absorption cross section of PSII according to previously published methods (Kolber et al., 1998; Oxborough et al., 2012). Excitation standards for each LED color were measured and each fluorescence trace normalized to its corresponding standard. Resulting fluorescence induction and relaxation curves were processed in R with the nonlinear curve (nlc) fitting package using equations adapted from (Kolber et al., 1998). Relaxation kinetics were fit to a double exponential decay equation (Suggett et al., 2022).

### Chlorophyll fluorescence-based phenotyping protocol

Each coral fragment was dark acclimated for 20-25 minutes and then phenotypic data were recorded using our custom chlorophyll a fluorometer and a 11-minute variable light protocol. For each measurement, excitation and relaxation protocols were sequentially repeated five times and averages for each wavelength recorded. This multispectral excitation and relaxation protocol was itself repeated 34 times in concert with an actinic light protocol which collected chlorophyll fluorescence data from coral samples during an initial dark period followed by three different light intensities (300, 50, 600 µmol m^-2^ sec^-1^) and a final dark recovery period (Fig. 1D). Spacing between each induction/relaxation was 6 seconds during the initial dark acclimated measurement (Kolber et al., 1998), and then 200 ms afterwards.

**Figure 1:**
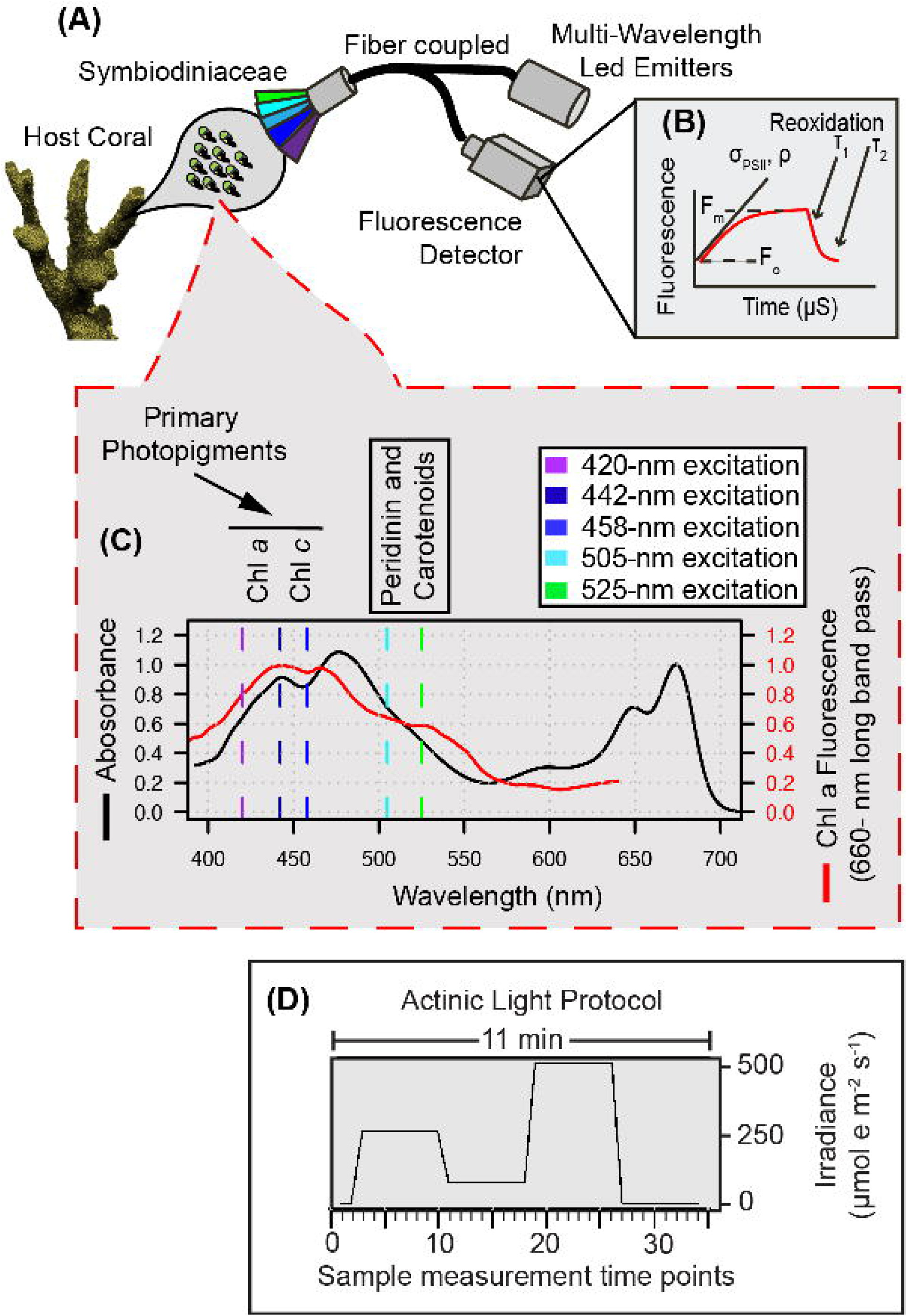
Multispectral conceptual illustration. Drawing of coral and symbiotic algae **(A)** illustrates the capture of a fluorescent signature where specific photopigments are preferentially excited by multispectral LEDs to resolve a µs fluorescence signal **(B)** captured through our single turnover system. Light absorption (black line) and fluorescence emission (red line) spectra from a representative algal culture **(C)** reflect differences in photopigment energy transfer that can be captured through multispectral excitation. Our benchtop fiber optic-based instrument exposes coral samples to an 11-minute protocol **(D)** during which multiple sampling events are carried out and enable the capture of spectrally dependent photosynthetic signatures in response to acclimation to three different irradiance levels and dark recovery.

The fluorescence induction and relaxation data measured with each excitation wavelength and at each step within the protocol, were used to calculate spectrally dependent maximum PSII quantum yields (Fv/Fm^ST^), PSII functional absorption cross-section (σ_PSII_), connectivity between reaction centers (ρ) and the two-time constants for photosynthetic electron transport (τ_1_ and τ_2_). While both time constants are derived from single turnover relaxation kinetics, τ_1_ reflects changes on the acceptor side of PSII, while τ_2_ better reflects changes in the time constant for plastoquinone pool reoxidation (Suggett et al., 2022). Additional data derived from our analysis include: non-photochemical quenching (NPQ ^ex-wavelength^) which was calculated as:

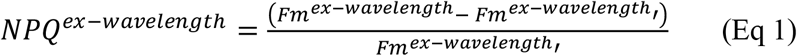

Where *Fm*^*wavelenghth*^ is the maximum fluorescence value recorded across all sample timepoints within the protocol. The maximum fluorescence value at each timepoint is represented by *Fm*^*ex*−*wavelenghth*^′ (Hennige et al., 2009). Photochemical quenching of chlorophyll *a* fluorescence (qP ^ex-wavelength^) was calculated as:

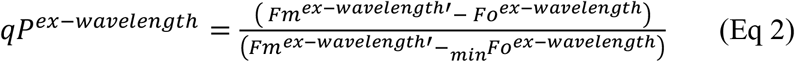

Here, _*min*_*Fo*^*ex*−*wavelenghth*^ is the minimum fluorescence value recorded across all sample timepoints within the protocol. For all samples, this occurred just after transition back into dark conditions. Similar to using far red light (van Kooten & Snel, 1990; Oxborough & Baker, 1997), the transition from high-light to dark leaves the plastoquinone pool in the highly oxidized state necessary for calculating _*min*_*Fo*^*ex*−*wavelenghth*^. Minimum fluorescence value at each timepoint is represented by *Fo*^*ex*−*wavelenghth*^ (Hennige et al., 2008; Hennige et al., 2011). Lastly, antennae bed quenching (ABQ^ex-wavelength^) was calculated as:

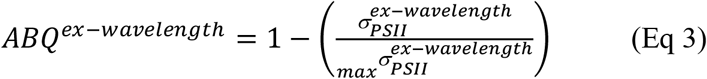

For ABQ calculations,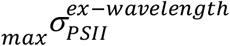 represents the maximum 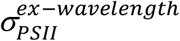 value across all sample timepoints within the protocol (Gorbunov et al., 2001; Hennige et al., 2009). Measurements for NPQ, qP and ABQ were each calculated independently for every excitation wavelength.

### Functional diversity sampling

Mote’s International Center for Coral Reef Research and Restoration (MML-IC2R3) on Summerland Key, Florida houses tens of thousands of coral fragments within their land-based nursery. The nursery contains approximately 60 land-based raceways (100” x 40” x 12”), which are supplied with filtered, UV sterilized, temperature controlled, near-shore seawater and maintained underneath 60% shade cloth canopies utilizing corrugated clear-plastic rain-guards as needed. Peak midday irradiance within these outdoor raceway aquaria was measured (Walz, 4pi sensor) at ∼400 µmol m^-2^ sec^-1^ under full sunlight (cloudless conditions). Overall, these systems represent highly similar light, temperature, and water movement conditions, and thus provide ideal conditions for assessing functional differences across endosymbiont species. For five of the six coral species from the outdoor aquaria (*Acropora palmata, Pseudodiploria strigosa, Siderastrea siderea, Pseudodiploria clivosa*, and *Orbicella faveolata*), between 9-18 individual fragments reflecting between 5-10 different genotypes were measured for algal phenotype and genotype. For each coral species, most genotypes were represented by a single coral fragment. However, between three and five fragments species^-1^ originated from a single genotype and were included to assess phenotypic resolution of the instrument. For *Stephanocoenia intersepta* only two different genotypes were measured (5 replicate fragments from one genet, and one frag of a second genet). A seventh species (*Montastraea cavernosa*), was also included in the study, but fragments originated from an indoor raceway aquaria on a 10:14 hr dark:light cycle utilizing LED lighting (175 µmol m^-2^ sec^-1^). Individual coral genotypes were pulled from Mote’s restoration broodstock, with individual fragments mounted to ceramic disks (Boston Aqua Farms, 3 cm diameter) and attached with cyanoacrylate gel (Bulk Reef Supply). All coral fragments utilized in this study had been acclimating to their respective environments for at least 2 months prior to our phenotype/genotype analyses.

### Genera level Symbiodiniaceae relative abundance determination

A small tissue sample was removed from each coral fragment and immersed in 2-ml of DMSO buffer and stored at 4°C. DNA was then extracted from preserved tissue samples using the Wizard extraction kit (Promega) and following standard protocols. An initial bead beating (1 minute using 0.5mm glass beads with tissue sample immersed in nuclei lysis solution) was included to aid in cell lysis. Quantity and quality of DNA was then assessed using a NanoDrop spectrometer (FischerSci). Only DNA with 260:230 and 260:280 ratios above 1 were utilized in downstream analysis. All samples were diluted to 2-ng ul^-1^ prior to qPCR. Next, methods similar to (Mieog et al., 2007; Mieog et al., 2009) were utilized to quantify the relative abundance of Symbiodiniaceae at the genera level within each sample. Genera specific primers targeting the Actin sequence were utilized to quantify relative abundance (McGinley, 2012; Grottoli et al., 2014). Copy number differences across genera were corrected for by utilizing genera specific DNA ladders derived from extracting DNA from a set number of Symbiodiniaceae cells. Cell cultures were utilized for *Cladocopium* (CCMP2466: NCMA), *Breviolum* (MF105b: originally isolated by M.A Coffroth), and *Symbiodinium* (CCMP2459: NCMA) whereas *Durusdinium trenchii* cells were separated and purified from the host coral *Montipora capitata* (symbiont identification confirmed via ITS2 sequencing and the Symportal database (Hume et al., 2019).Cell enumeration prior to each DNA extraction was completed using an Attune Nxt Flow Cytometer (Thermo Fisher Scientific). All primers were ordered from IDT-DNA and qPCR assays were performed using the PowerUp SYBR green master mix (Thermo Fisher Scientific) using a two-step protocol on an IQ5 quantitative PCR machine (Thermo Fisher Scientific) and according to previously established protocols for estimating relative abundance of Symbiodinaceae (Ulstrup & Van, 2003; LaJeunesse et al., 2009).

### Thermal Bleaching Experiment

Using the Climate and Acidification Ocean Simulator (CAOS) system located at MML-IC2R3, a two-month-long thermal bleaching experiment was performed on two branching coral species, *Acropora cervicornis* (n=10 genotypes) and *Acropora palmata* (n=5 genotypes). All coral fragments utilized in this study were acclimated for one month within the CAOS system prior to the start of the experimental treatment. Control and heat treatments consisted of a single shallow raceway per treatment, each with 10, 5-gallon flow through glass aquarium tanks (16” x 8” x 10”). One replicate fragment per genotype was distributed in each treatment tank, keeping coral species separate to avoid potential symbiont switching/shuffling. Filtered seawater was continuously supplied to each tank (14 L hr^-1^), with additional circulation provided by submersible pumps (120 gph, Dwyer). All experimental systems were maintained underneath 60% shade cloth canopies and clear plastic rain guards and experienced similar light levels as described above. At the start of the experiment (March 15^th^, 2021), the high-temperature raceway increased by 0.5 °C each day until a maximum of 30.5°C was reached. After one month of exposure to 30.5°C, the high-temperature raceway was increased another 1°C and remained at 31.5°C for an additional month. The ambient temperature raceway remained at 27 °C throughout the 2-month experiment (fig. S2).

Chlorophyl a fluorescence profiles (as described above) and reflectance measurements were measured on experimental days 64-65 (May 17^th^ and 18^th^ 2021 for *A. cervicornis* and *A. palmata* respectively). Fluorescence profiles were only collected from control coral fragments using a slightly modified protocol as the one described above. In this case, a total of 47 measurements were taken over a 16-minute time frame (initial dark period followed by three different light intensities: 500, 100, 750 µmol m^-2^ sec^-1^ and a final dark recovery period, see fig. S5a). For high-temperature treatment corals, only the initial dark-acclimated measurement (using 442nm excitation) was performed to derive (ΦPSII, σ_PSII,_ τ_1_,τ_2_). Absorption-based measurements were calculated on all coral fragments and achieved by measuring the reflectance spectra according to previously established methods (Rodriguez-Román et al., 2006). Briefly, our setup consisted of a white led (Luxeon) coupled to a USB2000 spectrophotometer (Ocean Optics) through a fiber-optic probe (Berkshire Photonics). Changes in absorbance can serve as a non-invasive proxy for monitoring changes in cell density/chlorophyll *a* content associated with coral bleaching (Rodriguez-Román et al., 2006; Hoadley et al., 2016). Here, absorbance readings at 420, 442, 458, 505 and 525-nm were selected to correspond with fluorescence excitation while an additional reading of 679nm reflects the maximum chlorophyll-*a* absorbance band. Bleaching response metrics for each coral genet are derived from absorbance and dark acclimated photo physiology (ΦPSII, σ_PSII,_ τ_1_,τ_2_ – calculated under 442nm excitation) and calculated as the % change between control and high temperature treatment measurements.

### Bleaching model generation and statistical analysis

All analyses were conducted in R (v.3.5.1) (Team, 2017). For photo-physiological comparisons, significant differences across algal genera were determined using a one-way ANOVA or Kruskal-Wallis if data did not fit assumptions of normality. Resulting datasets were then normalized using z-scores and 1000 bootstrap iterations were used to generate a heatmap and dendrogram using the R packages pvclust (Suzuki & Shimodaira, 2013) and dendextend (Galili, 2015). Custom scripts were utilized to identify the dominate photo-physiological metric within each heatmap row cluster (Fig. 2A). Average response profiles for each spectrally dependent ΦPSII, σ_PSII,_ τ_1_, and τ_2_ metric are reflected in Fig 3. For each spectrally dependent metric, a repeated measures linear mixed model with a Tukey posthoc (with Bonferroni correction) identified significant differences between the five selected phenotypes (algal/coral combinations) using the lmerTest (Kuznetsova et al., 2017) and multicomp (Hothorn et al., 2008) packages.

**Figure 2:**
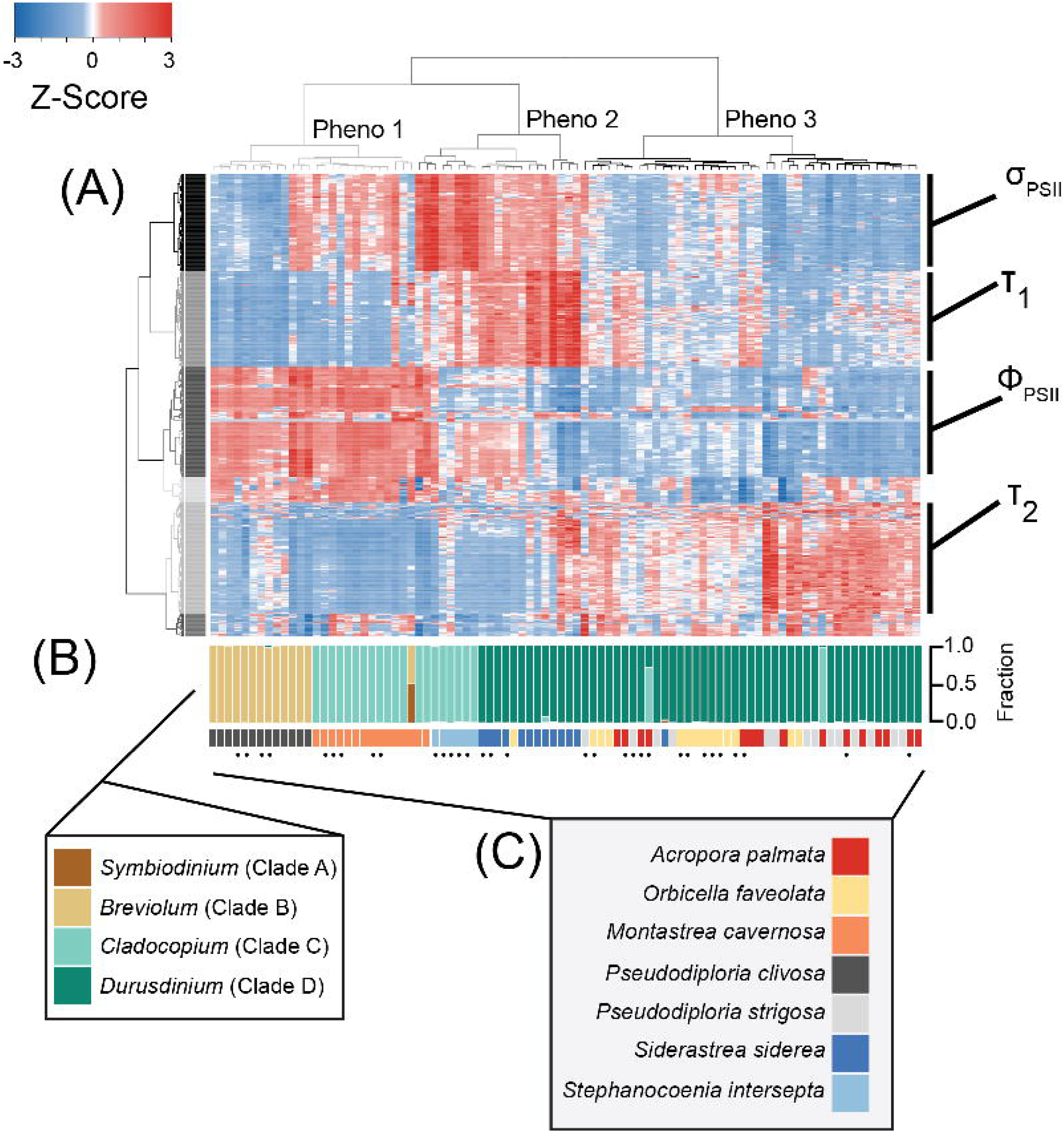
Phenotype to clade level comparison. Heatmap **(A)** reflects photo-physiological metrics collected using our multispectral and single-turnover instrument. Of the 1360 metrics measured, the heatmap represents only the 594 significantly different metrics across symbiont genera. Each row reflects a separate algal biometric displayed as standard deviation from the mean (z-score). Sample order was determined through hierarchical clustering reflected through the column dendrogram which identifies three distinct phenotypes (grey=1, dark grey=2, and black=3). Row order was also determined through hierarchical clustering generating 6 separate clusters indicated by colors. Dominant photo-physiological metrics within selected row clusters (four largest clusters) are indicated to the right of the heatmap. qPCR-based estimates for the proportion of Symbiodiniaceae genera within each sample are reflected in the upper barplot **(B)**. Host coral species is reflected in the lower barplot **(C)**. Replicate fragments from a single genet (within each species) are indicated by black dots.

**Figure 3:**
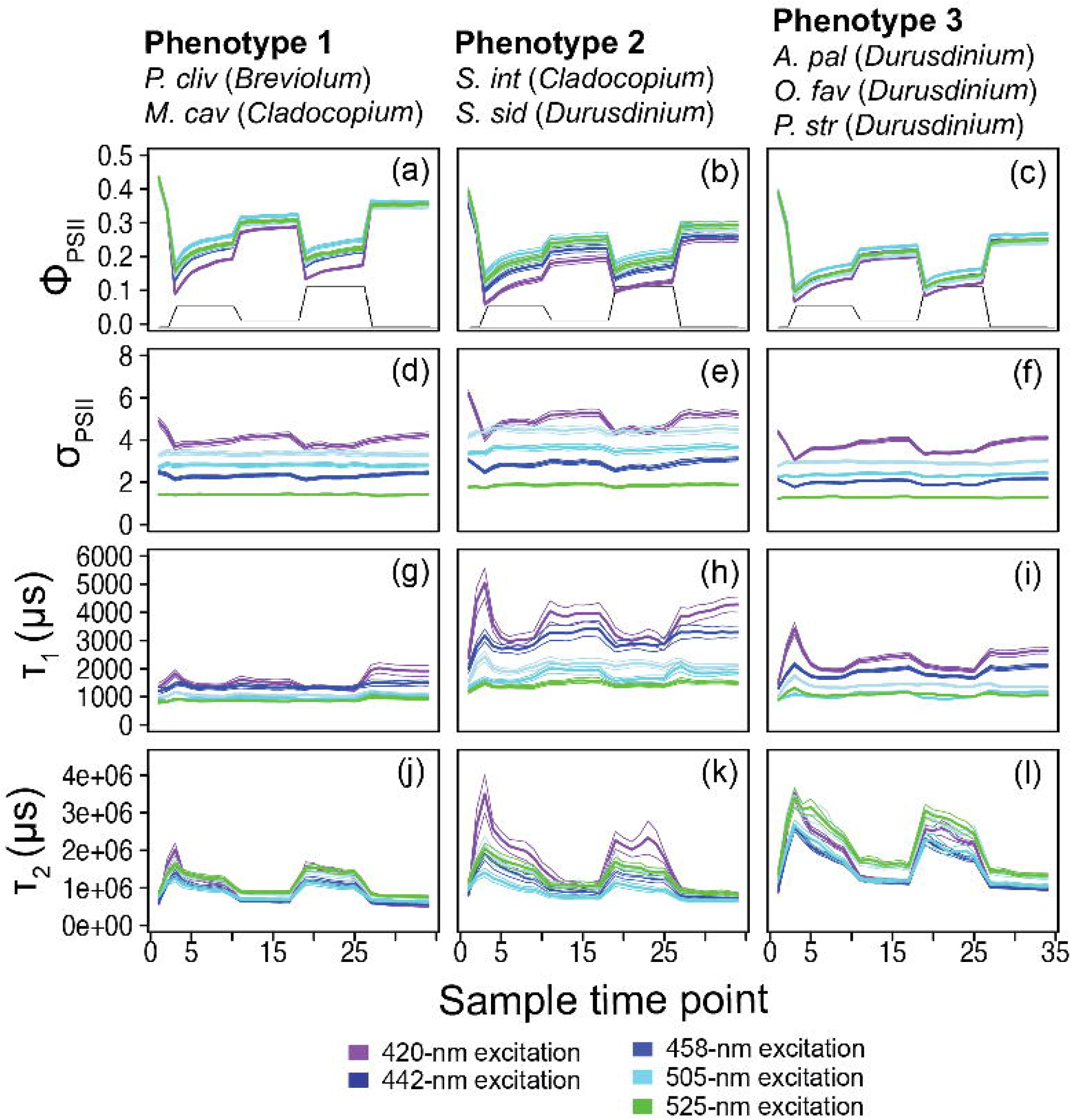
Phenotype profiles. Average (± standard error) traces for photo-physiological metrics identified within figure 2; quantum yield of PSII (Φ_PSII_; **A-C)** absorption cross-section of PSII (σ_PSII_; **D-F)**, and Tau_1_ (T_1_; **G-I)** and Tau_2_ (T_2_; **J-L)** reoxidation kinetics are displayed for phenotype 1(left column), phenotype 2 (middle column), and phenotype 3 (right column). Spectrally dependent responses for each of the five excitation wavelengths are reflected in each panel (thick bar = average, thin bars = standard error). For reference, the light protocol profile is inlaid within panels A-C.

In order to create the bleaching response model (Fig. 4), specific correlations (Pearsons value of 0.5 or above) between individual bleaching response metrics and fluorescence-based phenotyping data were identified using a network analysis (Fig. 4A) using the igraph package (Csardi & Nepusz, 2006). For each correlation, linear regression was used to first calculate the slope and then create a biometric value range associated with a bleaching response of less than ± 30% (Fig. 4B-C). Biometric value ranges were identified for each correlation and cumulatively represent our predictive bleaching model. To then test the accuracy of the model, individual values from coral fragment were scored according to if they were found to be within the established value range. The value output by the bleaching response model reflects the cumulative sum of scores for each fragment. Because the training data used to make our predictive model were created using a different protocol (47 sampling time points over 15 minutes) than the one used for characterizing the 7 coral species (34 sampling time points over 11 minutes) we used a truncated version of the protocol which selected data points that reflected similar conditions within the two protocols. Specifically, dark acclimated and transitional timepoints were used, while timepoints reflecting acclimation to light conditions were removed (fig. S5). A Shapiro-Wilks test was then used to confirm normal distribution in model output scores and significant differences across symbiont genera were then tested using a one-way ANOVA with a Tukey-posthoc. All algal biometric, bleaching response metrics, and qPCR data along with analytical scripts for generating figures 2-5 are available on GitHub (khoadley/BioOpticalBleachingPrediction2022).

**Figure 4:**
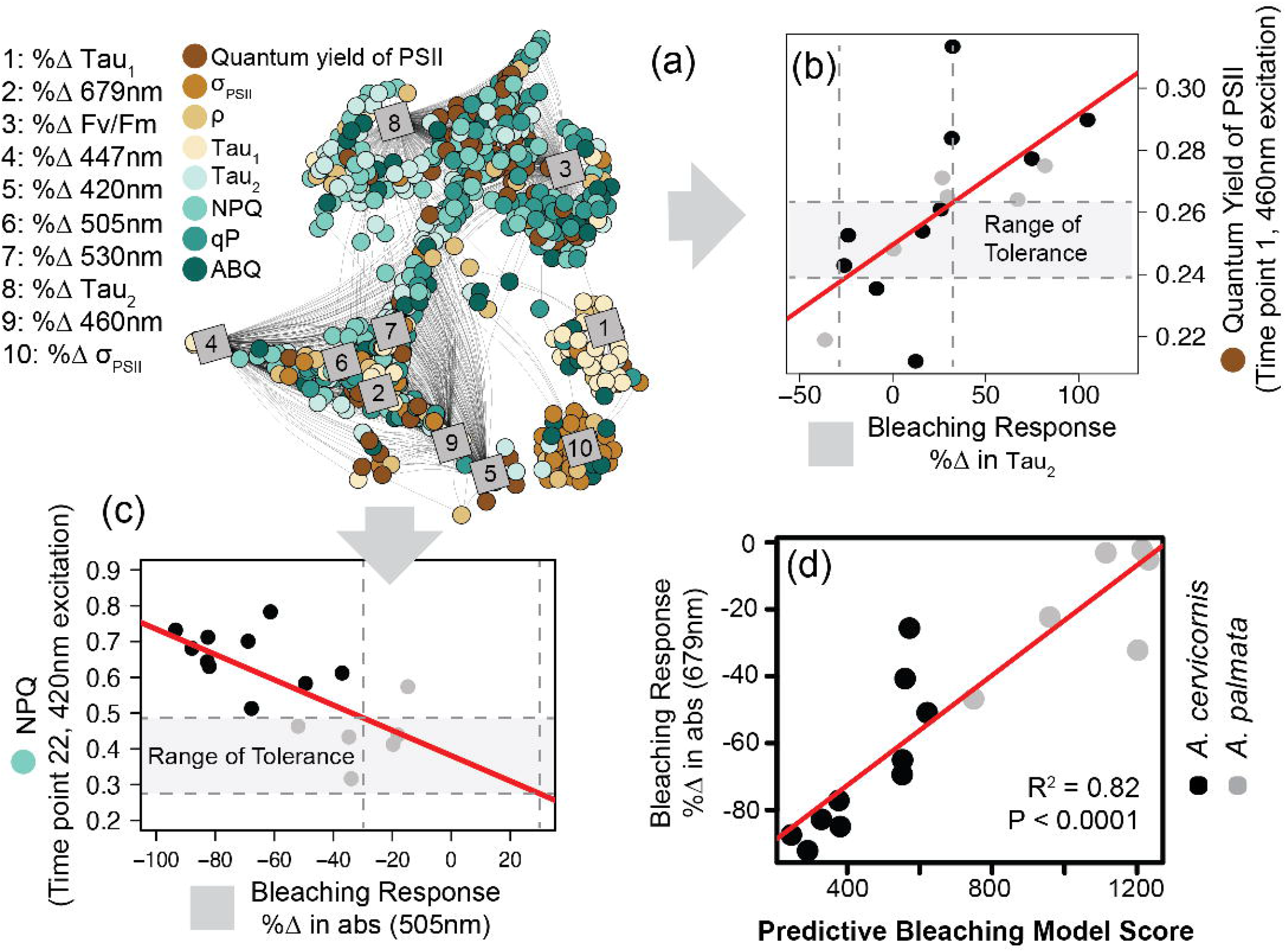
Model training and predictive bleaching output. Network analysis **(A)** reveals strong correlations between specific algal biometrics (circles; measured by our novel instrument and protocol) with coral bleaching response metrics (squares; measured as % change in response to high temperature). Connecting lines reflect significant correlation (Pearson > 0.5). Representative correlations pulled from the analysis show a strong relationship between the quantum yield of PSII (Φ_PSII_^460^) and the % Δ in Tau_2_ **(B)** or light energy dissipation (NPQ^420^) and the % Δ in absorbance at 505nm **(C)**. A range of tolerance is then constructed for each algal photo-physiological-bleaching response metric combination identified in the network analysis. Vertical dashed lines outline a ± 30% change in response to thermal stress while horizontal dashed lines outline the range of tolerance (gray outline). The resulting model was then tested against the training data by ranking each coral genotype based on the number of photo-physiological metrics that fall within their corresponding range of tolerance (higher model numbers suggest greater thermal tolerance).**(D)** The accuracy of the predictive bleaching model was compared to a coral bleaching proxy (% Δ in absorbance at 679nm) and the strength of the correlation was tested using linear regression (red lines).

## Results

### Multispectral approach and protocol

A total of 8 (Table 1) different photo-physiological parameters can be measured (Fig. 1B) or calculated (equations 1-3) from chlorophyll-*a* fluorescence derived from each separate LED color utilized (420, 442, 458, 505, 525nm – Fig. 1C). For each sample, an individual measurement timepoint (Fig. 1D) generates a total of 40 different algal biometrics (5 LED colors * 8 photo-physiological parameters). Between 34 (seven coral species) and 47 (thermal bleaching assay) measurement timepoints were utilized to characterize photo-physiological response during the actinic light protocol (Fig. 1D and fig. S4a) and generated 1360 (40 algal biometrics * 34 measurement timepoints) and 1880 (40 algal biometrics * 47 measurement timepoints) individual algal biometrics respectively. These ‘high-content’ datasets contain both subtle and dynamic differences in how each sample responds to variable light conditions and are collectively utilized to establish phenotypes and predictive bleaching models as described below.

### Phenotype to genera profiles

Quantitative PCR was performed to estimate the relative abundance of the four most common symbiont genera: *Symbiodinium* (formerly clade A), *Breviolum* (formerly clade B), *Cladocopium* (formerly clade C) and *Durusdinium* (formerly clade D) within each coral fragment. Two of the eleven *A. palmata* genotypes tested contained predominantly (>95%) *Cladocopium* symbionts whereas the other nine genotypes contained predominantly *Durusdinium* symbionts. All *O. faveolata, S. siderea* and *P. strigosa* coral fragments also contained predominantly *Durusdinium* symbionts. A single fragment of *M. cavernosa* contained an even split between *Symbiodinium* and *Breviolum* symbionts whereas all others were dominated by *Cladocopium*. All *P. clivosa* coral fragments were dominated by *Breviolum* symbionts. Lastly, all six fragments (5 ramets of one genotype and one of a second genotype) of *S. intersepta* were dominated by *Cladocopium* symbionts (Fig. 2B).

Of the 1360 fluorescence-based algal biometrics derived from the actinic light protocol used to characterize our seven coral species, 594 differed significantly across the symbiont genera found across our samples. These significant algal biometrics were then utilized to create a dendrogram which organized coral fragments into three major phenotypes based on the largest clustering groups (Fig. 2A). All *P. clivosa* (*Breviolum*) and, except for two fragments, all *M. cavernosa* (*Cladocopium*) clustered together to form a single phenotype (phenotype 1).

*Durusdinium sp*. symbionts within the coral *S. siderea* (along with a single fragment of *A*. palmata) and two *Durusdinium* dominated *M. cavernosa* coral fragments clustered together to form a second phenotype (phenotype 2). Except for a single fragment of *O. faveolata*, all other *Durusdinium* dominated corals (*O. faveolata, A. palmata* and *P. strigosa*) formed a final cluster (phenotype 3) which included two *A. palmata* fragments with *Cladocopium* (Fig. 2A). The quantum yield of PSII (ΦPSII), absorption cross-section of PSII (σ_PSII_) and T_1_ and T_2_ reoxidation kinetics were identified as being the most important photo-physiological parameters in creating the observed phenotypic structure across coral fragments. Spectrally dependent profiles for these four photo-physiological parameters were therefore plotted in figure 3 and reflect how each of our three distinct phenotypes responded throughout all measurement timepoints during the actinic light protocol.

For ΦPSII^420^, ΦPSII^442^ and ΦPSII^460^ profiles, all three phenotypes were significantly (*p* < *0.0001*) different with phenotype 1 exhibiting the highest and phenotype 3 the lowest values. However, phenotypes 2 and 3 had similar ΦPSII^505^ and ΦPSII^525^ profiles which were significantly (*p* < *0.0001*) lower than those observed for phenotype 1 (Fig. 3A-C). Significantly (*p* < *0.0001*) higher σ_PSII_^420^ values were observed for phenotype 2 as compared to both phenotype 1 and 3. The remaining four spectrally dependent σ_PSII_ profiles also had significantly (*p* < 0.05) different profiles with the highest and lowest values found in phenotypes 2 and 3 respectively (Fig. 3D-F). For all five spectrally dependent τ_1_ reoxidation profiles, phenotype 2 produced significantly (*p* < *0.0001*) higher values as compared to phenotypes 1 and 3. The lowest values for τ_1_ were almost always observed in phenotype 1 except for τ ^505^ where phenotype 3 also contained similarly low values across the profile (Fig. 3G-I). Lastly, τ_2_^420^ reoxidation kinetic profiles were significantly (*p* < *0.001*) higher in phenotypes 2 and 3 as compared to 1. However,τ_2_^442^ profiles differed significantly across all three profiles (*p* <*0.x001*) whereas τ_2_^460^, τ_2_^505^, and τ ^525^ profiles were significantly lower in both phenotypes 1 and 2 as compared to phenotype 3 (Fig. 3J-L).

### Bleaching response model development and training

For each control fragment of *A. palmata* and *A. cervicornis* utilized in the thermal bleaching experiment, a total of 1880 algal biometrics were derived from our fluorescence-based phenotyping protocol. For each genotype, absorbance-based measurements from control and experimental fragments were also obtained to quantify the % change in absorbance calculated for six specific wavelengths which correspond to the five fluorescence excitation peaks (420, 442, 458, 505, and 525nm) plus the chlorophyll-*a* absorption maxima (679nm). Additionally, dark-adapted measurements of the quantum yield of PSII (ΦPSII^442^), the absorption cross-section of PSII (σPSII^442^), τ_1_442 and τ_2_^442^ reoxidation kinetics were also obtained for each genotype’s control and treatment fragments and are similarly represented as the % change in response to thermal stress (fig. S4). These 10 bleaching response metrics (absorbance + dark adapted photo-physiological metrics) were then tested for correlation with all 1880 algal biometrics using a network analysis. Of the 18,800 total correlations possible, 1973 were identified as having a Pearson value of 0.5 or above (Fig. 4A). These identified biomarkers of thermal stress were then utilized to score individual coral genotypes according to the number of values that were contained within each corresponding ‘range of tolerance’. The bleaching response model number reflects the cumulative sum of scores and the accuracy of this score was assessed by plotting model number against the % change in absorbance at 679nm. This absorbance metric was chosen as it best reflects changes in absorbance that are related to reductions in symbiont cell density or chlorophyll content within the host tissue (Rodriguez-Román et al., 2006; Hoadley et al., 2016). A linear regression between the model scores and % change in absorbance values indicate high correlation (R^2^=0.82, *p* < *0.0001*) and thus good predictive power (Fig. 4D).

### Application of bleaching prediction model across seven coral species

A truncated version (fig. S4) of the predictive bleaching model was further tested on the seven coral species for which the phenotype to genera profiles were characterized (Fig. 2). Of the 1973 different algal biometric/bleaching response metric correlations identified from the *Acropora* bleaching data, only 259 algal biometrics were utilized in the truncated version (Fig. 5) and represent similar light conditions across our two different protocols (fig. S4). Despite using a smaller dataset to predict bleaching sensitivity, model scores for *Durusdinium* were significantly (*P* < *0.001*) higher than those for *Breviolum* and *Cladocopium* dominated coral fragments (Fig 5).

**Figure 5:**
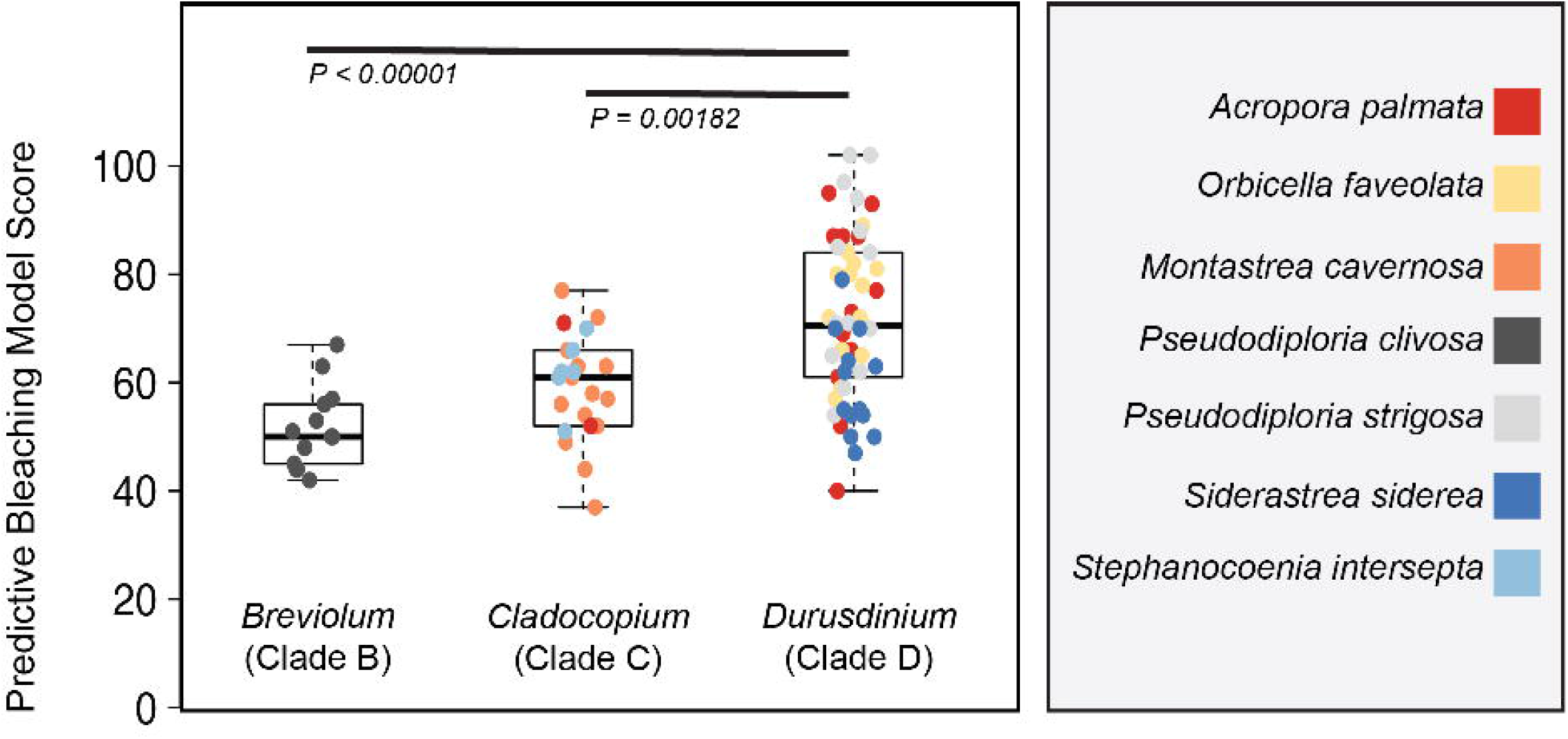
Model output on clade level comparisons. A reduced model containing only 261 of the original 1973 comparisons was applied to photo physiological data from each of the 90 coral fragments represented in figure 3. Box plots represent the mean and ± 1 standard deviation of model output for *Symbiodinium, Breviolum* and *Durusdinium-*dominated corals. Significant differences between symbiont types are indicated by black bars. Individual coral fragments are also plotted with color reflecting coral species.

## Discussion

Our predictive model takes advantage of the cumulative power derived from the incorporation of many photo-physiologically based biomarkers of thermal stress. For many corals, tolerance to environmental perturbations such as high temperatures is directly linked to symbiont species and our instrumentation and analytical pipeline directly targets underlying photo-physiological differences between species. Phenomic techniques that incorporate ‘high-content’ physiological datasets and multi-variate analyses have previously led to critical breakthroughs in agricultural fields (Furbank & Tester, 2011; Furbank et al., 2019) and a similar approach could prove beneficial within both basic and applied fields of coral reef science. Fluorescence-based tools are gaining popularity as a means of non-destructively generating photosynthetically-based phenotypes of symbiotic algae (Voolstra et al., 2020; Cunning et al., 2021; Suggett et al., 2022) that can also produce direct, actionable results. While our model output did assign higher bleaching scores to corals dominated by symbionts with well characterized thermal tolerance (Fabricius et al., 2004; LaJeunesse et al., 2009; Ortiz et al., 2012), additional direct testing of predictive model scores is still needed. Nevertheless, new tools and analytical pipelines that help bridge the gap between current and future coral conservation and restoration techniques are urgently needed (Vardi et al., 2021; Voolstra et al., 2021a) and highly scalable methods such as those utilized here could provide such an avenue.

### Phenotype to clade level comparisons

While the case for chlorophyll-*a* based phenotyping is increasing in popularity (Hennige et al., 2009; Hoadley et al., 2019; Hoadley et al., 2021; Suggett et al., 2022), the current study expands on this further by using a novel system and protocol to massively increase the number of individual algal biometrics collected from each sample. Combined with a global analysis of the resulting data, our work directly connects phenotypic differences with either host species or Symbiodiniaceae clade-level estimates of genetic variance from 90 coral fragments (Fig. 2). Of the four photo-physiological parameters identified to be most valuable in establishing distinct phenotypes (Fig. 2A), only the quantum yield of PSII can be measured (albeit at a single excitation wavelength) using multi-turnover fluorometers such as the commonly used and commercially available PAM (Walz). The remaining three photo-physiological parameters (σ_PSII_, τ_1_ and τ_2_) can only be derived from single turnover instrumentation (Suggett et al., 2022). Overall, measurements of τ_1_ and τ_2_ indicate slower electron transport rates through the photochemical apparatus in *Durusdinium* phenotypes while the more dynamic responses to changing light observed for these two parameters may also be indicative of differences in how downstream photochemical activity is regulated across Symbiodiniaceae genera or species (Roberty et al., 2014; Vega de Luna et al., 2020). The high level of algal biometric parameterization demonstrated in our approach is valuable for identifying phenotypic differences. In addition, the use of multi-spectral excitation allows for even further non-destructive interrogation that incorporates differences in photopigment utilization across species and/or environments.

The absorption cross-section of PSII (σ_PSII_) measures the proportion of light captured by light-harvesting compounds (LHCs) and spectrally resolved differences in this parameter are particularly well suited for understanding how different photopigments are utilized under various conditions (Szabó et al., 2014; Hoadley & Warner, 2017). The family Symbiodiniaceae contains several different photopigments, some are involved with light-harvesting while others function as accessory pigments which either channel additional energy towards the photosynthetic reaction centers or dissipate excess light energy as fluorescence and heat (Iglesias-Prieto & Trench, 1994; Iglesias-Prieto & Trench, 1997). Large changes in 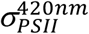 and 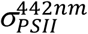 values are observed during the actinic light protocol (Fig. 3D-F) and likely indicate changes in the orientation of chlorophyll-a containing LHCs to downregulate the quantity of light energy transferred to PSII reaction centers (Iglesias-Prieto et al., 1991; Iglesias-Prieto et al., 1993; Niedzwiedzki et al., 2013; Niedzwiedzki et al., 2014). In contrast, smaller responses to changes in the actinic light protocol are observed for 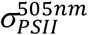 and 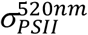 values which preferentially target peridinin containing LHCs and carotenoids (diadinoxanthins). The spectrally dependent differences in *σ*_*PSII*_ observed in response to changing light conditions showcase the utility of using a multispectral approach to delineate photopigment specific variability. However, future research that directly links multispectral *σ*_*PSII*_ output to photopigment concentrations and connectivity to reaction centers is needed.

### Linking phenotype to thermal stress - an optical approach for bleaching prediction

Algal biometric data and bleaching response metrics collected from *A. cervicornis* and *A. palmata* coral colonies were utilized to identify photo-physiologically based biomarkers of thermal stress to train a predictive bleaching model (Fig. 4A-B) which then evaluated coral colony thermal tolerance based on algal biometric data alone. High accuracy in the model output (R^2^=0.82) was achieved when tested against the same coral fragments used for training (Fig. 4D), where *A. cervironis* colonies experienced greater thermal stress than *A. palmata*. Within the greater Caribbean, both *Acropora* species are often found in association with the symbiont, *Symbiodinium fitti* (LaJeunesse, 2002). However, under nursery conditions, many *A. palmata* fragments appear to have switched to the symbiont genera *Durusdinium* (Fig. 2B), which contains many species that are well documented as thermally tolerant (Fabricius et al., 2004; LaJeunesse et al., 2009; Ortiz et al., 2012). While not directly tested within our experimental coral colonies, presumed differences in symbiont species across the two *Acropora* coral species likely regulates the distinct phenotypes (fig. S3a) observed. The linkage between bleaching responses and fluorescence-derived phenotypes are what enables the rapid identification of hundreds of different biomarkers of thermal stress using our network analysis. For example, non-photochemical quenching (NPQ) values from *A. cervicornis* were typically higher than those from *A. palmata* colonies. High NPQ under ambient conditions may indicate that the coral colony is already operating at its environmental limit, making it particularly vulnerable to further stress such as a high-temperature event (Hennige et al., 2011). Indeed, NPQ values are linked to various bleaching response metrics (Fig. 4A). While many algal biometrics display reasonable correlations with bleaching response metrics (Pearson R values between 0.5 and 0.85) the cumulative effect of using hundreds of biomarkers is what provides the high model accuracy (Fig. 4D, Pearson R value = 0.92) when applied to the same coral colonies.

To further assess the use of algal biometrics to predict bleaching tolerance, we applied a truncated version of our model to the larger dataset of seven coral species for which we only collected algal biometric data. While accuracy of the model output cannot be directly assessed as these individual fragments were not thermally challenged, higher scores from corals dominated by *Durusdinium* suggest bleaching predictions follow our current understanding of thermal tolerance across symbiont genera (Fabricius et al., 2004; LaJeunesse et al., 2009; Ortiz et al., 2012; LaJeunesse et al., 2018). Importantly, these results were achieved using a truncated model and applied to samples measured using a different actinic light protocol (fig. S5). The application of more purpose-built models would no doubt improve results. Nevertheless, our results clearly show predictive ability based on the unique phenotypic profile of *Durusdinium* symbionts.

Interestingly, model output values for *S. siderea* coral colonies were more variable despite also being dominated by *Durusdinium* and may point to either population level differences in symbiont physiology or host-influence on symbiont phenotype. While prior studies found only minimal population structure for *Durusdinium* within the greater Caribbean region (Pettay et al., 2015), host-specific traits such as green fluorescence protein, tissue thickness, skeletal structure and nutrient regulation can alter the light and nutrient environment for their symbiotic algal partners (Rodriguez-Román et al., 2006; Enríquez et al., 2017; Wangpraseurt et al., 2017a; Wangpraseurt et al., 2017b; Xiang et al., 2020; Bollati et al., 2022). For *D. trenchii* symbionts harvested from inshore reef habitats in Palau, differences in cellular physiology were attributed to host coral species (Hoadley et al., 2019). While not directly attributable to the host, *S. siderea* coral fragments clustered separately from all other *Durusdinium-*dominated corals, indicating strong phenotypic differences across coral species potentially driven by host traits. Importantly, these phenotypic differences (Fig. 2A-C) likely also drive the lower model output for *S. siderea* as compared to other *Durusdinium-*dominated coral fragments. Although further research is needed, alterations to the symbiont environment driven by differences in coral species or even genotype might also be detected with our phenomic approach, providing a way to incorporate host-bleaching response mechanisms into our analytical framework.

### Future application towards trait-based selection of corals for restoration initiatives

Coral restoration strategies such as assisted evolution aim to improve resilience through propagation of corals with desirable traits including thermal tolerance (Anthony et al., 2017; Voolstra et al., 2021a). However, rapid and scalable identification of coral genotypes with desirable traits is challenging (Baums et al., 2019; Morikawa & Palumbi, 2019; Voolstra et al., 2021b) and most techniques require infrastructure and resources well outside the reach of many restoration facilities. For restoration initiatives focused on coral species where thermal stress is largely derived from the symbiont species, tools that capture functional trait differences could be beneficial in selecting thermally resilient colonies.

This study reflects a novel instrument platform and analytical pipeline, where results from a single experimental bleaching assay were applied towards predicting functional traits within a separate set of coral colonies. While our results were largely positive and reflect our current understanding of bleaching tolerance across symbiont genera, direct testing of predictive bleaching scores is still needed before this approach can be considered as a potential tool for scaling up coral restoration initiatives. If future studies indeed show good correlation between model score output and bleaching tolerance, our analytical pipeline could provide a scalable means for assessing thermal tolerance.

The utilization of multispectral CF techniques to rapidly predict bleaching susceptibility in reef corals by capturing the unique phenotypic characteristics of the symbiotic algae may hold promise for certain coral species and environmental conditions. However, traits which regulate thermal tolerance are highly variable across and within species and environments (Weis, 2008; Weis & Allemand, 2009; Barshis et al., 2013; Bay & Palumbi, 2014; Suggett et al., 2017). Consequently, the efficacy of approaches that utilize symbiont phenotypes for assessing thermal tolerance likely differ across coral species and environmental conditions.

Scaling-up the identification of thermally tolerant colonies will require breakthroughs in accessibility to modern techniques, and open-source tools such as our fluorometer could play a critical role in bridging this important gap. Our analytical pipeline collapses complex photo-physiological biomarkers into a cumulative score that can be easily utilized by scientists and restoration practitioners alike. While our study only represents a conceptual framework for the relatively instantaneous assessment of bleaching susceptibility, future work will need to directly test bleaching scores, along with the long-term efficacy of this approach on both nursery and out planted coral fragments.

## Acknowledgements

We thank the staff and interns at Mote’s International Center for Coral Reef Research and Restoration on Summerland Key, FL for their assistance and support. Research activities were made possible through the Florida Fish and Wildlife Conservation Commission (SAL-21-1724-SCRP and SAL-21-2048-SCRP and Florida Keys National Marine Sanctuary (FKNMS-2015-163-A3 and FKNMS-2017-136) permits. The work was funded by the National Science Foundation, grant nos. 2054885 to K.D. Hoadley and the State of Florida via the Florida Fish and Wildlife Conservation Commission contract #20151 to Mote Marine Laboratory. The authors declare no competing interests.

## Author Contributions

K.H. and G.L. developed all hardware and software associated with the prototype instrument. K.H. and E.M. planned and designed the research. K.H., A.M., B.P., S.L., S.W., and C.P. and CK. performed experiments, conducted fieldwork, and analyzed data; K.H. wrote the manuscript. All authors provided feedback on the manuscript. K.H. agrees to serve as the author responsible for contact and ensures communication.

## Data Availability Statement

All data needed to evaluate the conclusions in the paper are present in the paper and/or the Supplementary Materials. Pending scientific review, instrument development plans, software, and processing scripts, along with raw data and analytical scripts for Figures 3 - 6 will be available via github (khoadley/fluorometer_2022, and khoadley/Mote_2021).

