## supplemental figures for "A phenomic modeling approach for using chlorophyll-a fluorescence-based measurements on coral photosymbionts: a step towards bio-optical bleaching prediction"

**This PDF file includes:**

Supplementary Text
Figure S1
Figure S2
Figure S3
Figure S4

**Fluorometer build:** The multispectral fluorometer runs on a single PCB board which integrates excitation LED drivers, actinic light drivers, and analog input. Specific PCB blueprints along with individual components for populating the board are available on khoadley/MultiSpectral-ST-PCB. The PCB board is located underneath the optical bench along with power supplies for the PCB board and avalanche photodiode. Firmware for the teensy microcontroller is written using the Arduino IDE and a python-based software program communicates with the microcontroller and manages data capture. All firmware and software are open-source and available via GitHub (khoadley/MultiSpectral-ST-PCB). Each measurement creates two files (a raw data file and a preliminary processing file). Both are stored on the teensy micro-SD card until downloaded unto a computer (either using the python-based GUI, or directly from the micro-SD card). Raw CSV files are then processed using a custom R script, also available via GitHub.

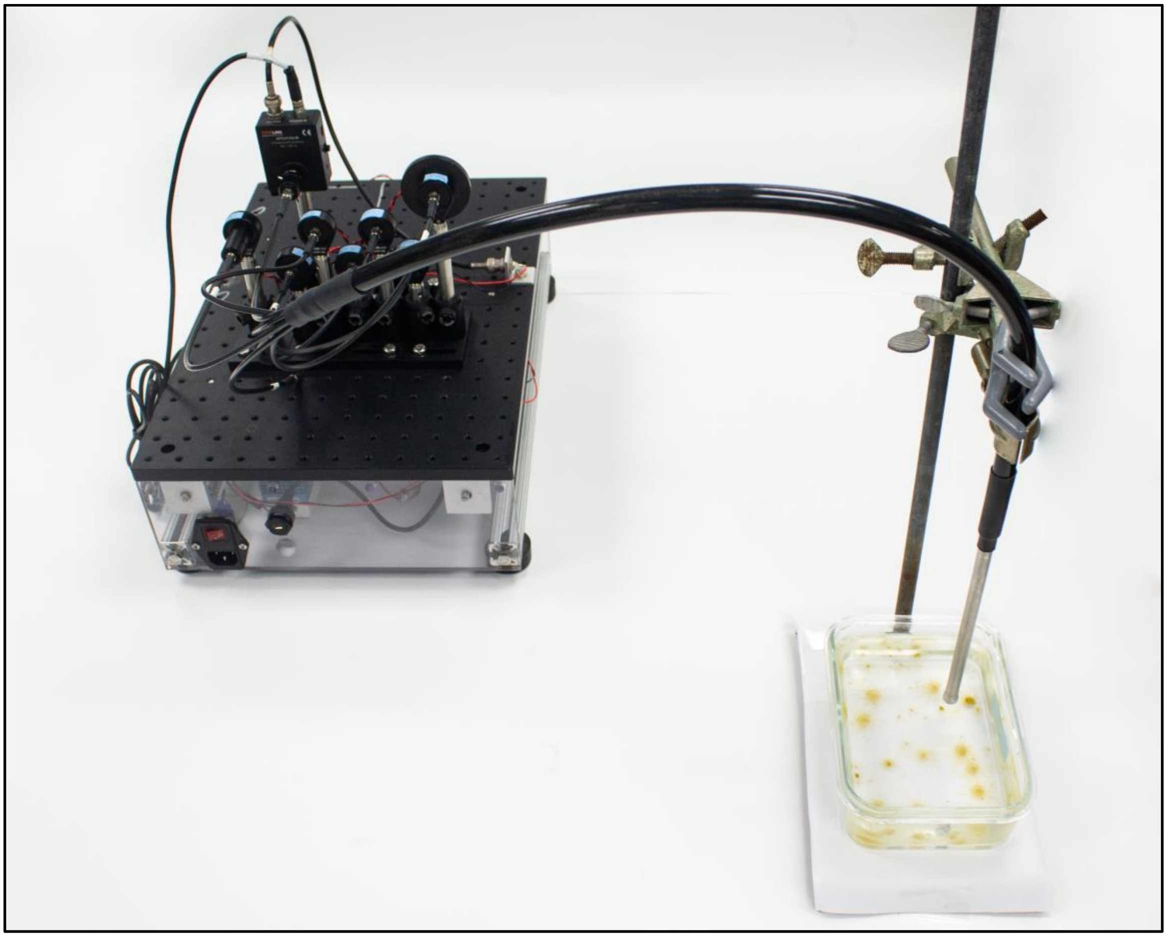

**Supplemental figure 1: Instrument and light protocol.** Multispectral excitation and capture of fluorescent signatures were carried out continuously using our benchtop and fiber optic-based instrument **(A)** which exposes a single coral sample to an 11-minute protocol, during which 34 sampling events are carried out. This protocol design captures spectrally dependent photosynthetic signatures in response to acclimation to three different light levels and dark recovery. Fiber optic cable couples LED and detector to a common end. As an example, the common end is submersed in a container with *Aiptasia*. Photo taken by Audrey McQuagge.

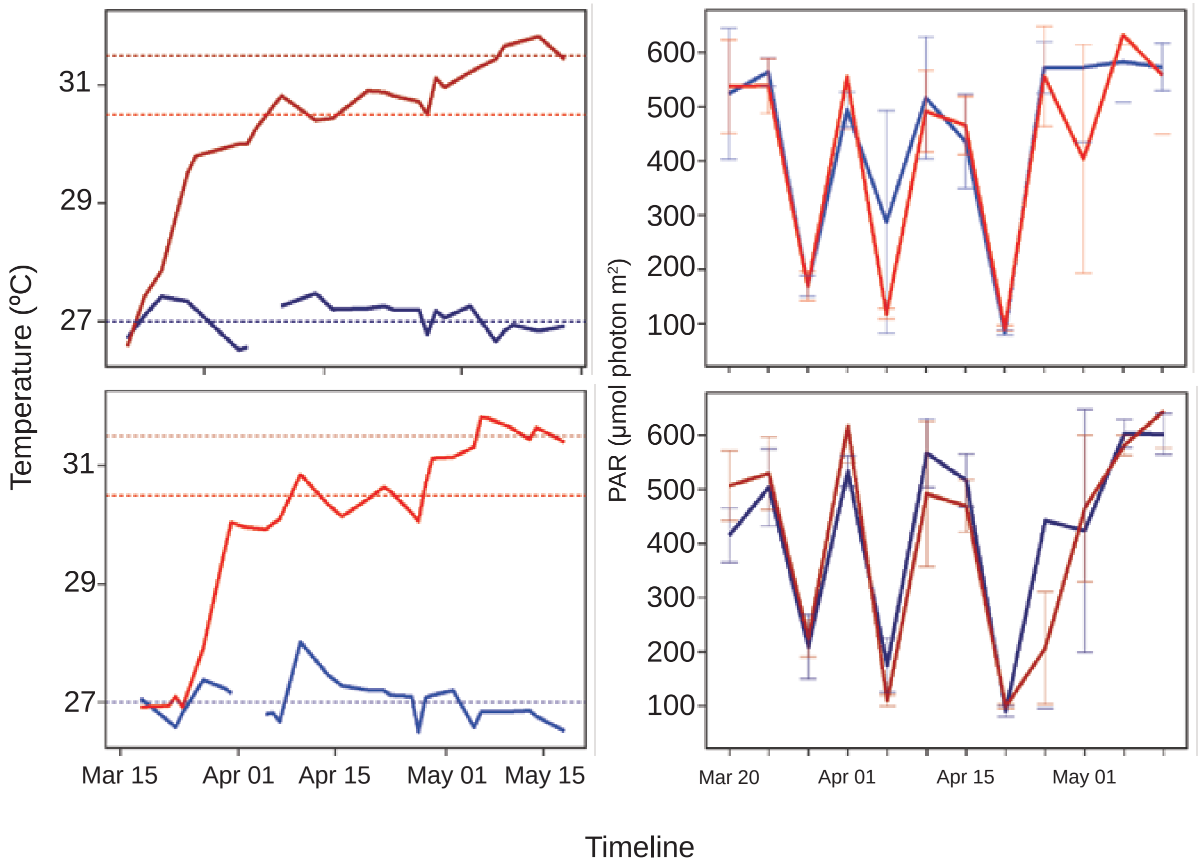


**Supplemental figure 2: Experimental bleaching conditions.** Temperature (left) and light (right) profiles for each treatment condition (control=blue, heat=red) display thermal ramping along with inherent daily variability in temperature and light throughout the full two-month experiment. Horizontal dashed lines represent target treatment temperatures. A single light measurement (PAR) was taken at midday every two or three days and reflects variability in midday light levels across experimental days, but similar irradiances across treatment tanks.

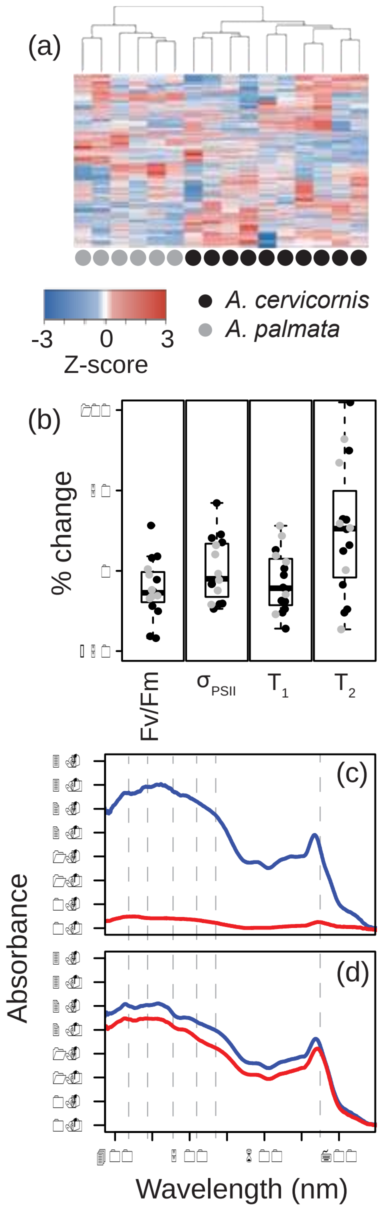


**Supplemental figure 3: Photo physiological data and bleaching response metrics for model input**. All photo physiological data derived from control coral fragments using our instrumentation are displayed in the heatmap **(a)** which appears to cluster samples across coral species. Box and whisker plots reflect the % change (from control and high temperature treatments) in maximum quantum yield (Fv/Fm), absorption cross section ($\sigma_{PSII}$), and Tau$_1$ and Tau$_2$ reoxidation kinetics. Individual genotype responses for *A. cervicornis* (black circles) and *A. palmata* (gray circles) are overlaid on each plot. Representative spectral absorbance profiles for a thermally susceptible **(c)**, and resilient **(d)** coral genotype. For each coral genotype, absorbance traces reflect fragments in control (blue line) and high temperature (red line) treatments. Dotted gray lines indicate the specific wavelengths where % change in response to high temperature was calculated and subsequently utilized to train the predictive bleaching model (Fig 5).

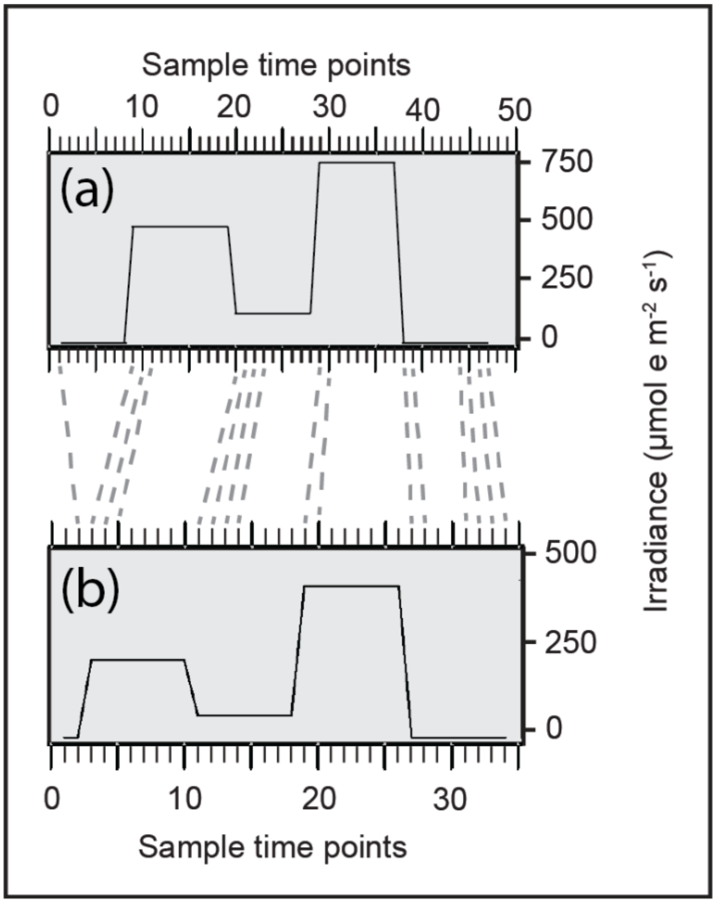


**Supplemental figure 4. Truncated predictive bleaching model:** Because the training data used to make our predictive model was created using a different actinic light protocol (**A** - 47 sampling time points over 15 minutes) than the one used for characterizing the 7 coral species (**B** - 34 sampling time points over 11 minutes) we used a truncated version of the protocol which selected data points that reflected similar conditions within the two protocols (grey dotted lines). Specifically, dark acclimated and transitional timepoints were used, while timepoints reflecting acclimation to light conditions were removed due to differences in light intensity. Only biometrics from these specific areas were utilized to construct our truncated model and test bleaching resilience within our 7 coral species data set.
